# The 5’ untranslated region of the *EFG1* transcript promotes its translation to regulate hyphal morphogenesis in *Candida albicans*

**DOI:** 10.1101/328948

**Authors:** Prashant R. Desai, Klaus Lengeler, Mario Kapitan, Silas Matthias Janßen, Paula Alepuz, Ilse D. Jacobsen, Joachim F. Ernst

## Abstract

Extensive 5’ untranslated regions (UTR) are a hallmark of transcripts determining hyphal morphogenesis in *Candida albicans.* The major transcripts of the *EFG1* gene, which are responsible for cellular morphogenesis and metabolism, contain a 5’ UTR of up to 1170 nt. Deletion analyses of the 5’ UTR revealed a 218 nt sequence that is required for production of the Efg1 protein and its functions in filamentation, without lowering the level and integrity of the *EFG1* transcript. Polysomal analyses revealed that the 218 nt 5’ UTR sequence is required for efficient translation of the Efg1 protein. Replacement of the *EFG1* ORF by the heterologous reporter gene *CaCBGluc* confirmed the positive regulatory importance of the identified 5’ UTR sequence. In contrast to other reported transcripts containing extensive 5’ UTR sequences, these results indicate the positive translational function of the 5’ UTR sequence in the *EFG1* transcript, which is observed in context of the native *EFG1* promoter. The results suggest that the 5’ UTR recruits regulatory factors, possibly during emergence of the native transcript, which aid in translation of the *EFG1* transcript.

**IMPORTANCE:** Many of the virulence traits that make *Candida albicans* an important human fungal pathogen are regulated on a transcriptional level. Here we report an important regulatory contribution of translation, which is exerted by the extensive 5’ untranslated regulatory sequence (5’ UTR) of the transcript for the protein Efg1, which determines growth, metabolism and filamentation in the fungus. Presence of the 5’ UTR is required for efficient translation of Efg1, to promote filamentation. Because transcripts for many relevant regulators contain extensive 5’ UTR sequences, it appears that virulence of *C. albicans* depends on the combination of transcriptional and translation regulatory mechanisms.

## Introduction

Transcriptional networks are known to govern growth and virulence of the human fungal pathogen *C. albicans.* Transcription factors have been identified that regulate the interconversion between its yeast cell form and a filamentous hyphal form, or the rod-like *opaque* form. Efg1 is a key bHLH-type regulatory protein that controls hyphal morphogenesis in a dual manner, promoting filamentation under normoxia in the presence of environmental cues (1, 2), but repressing it under hypoxia (3, 4). Its promoting function depends on increased histone acetylation and chromatin remodelling at promoters of target genes (5), which facilitates hyphal initiation; shortly thereafter, however, *EFG1* expression is strongly downregulated to prevent its interference with subsequent processes required for hyphal formation (6, 7). Under hypoxia, Efg1 represses the expression of genes encoding hyphal inducers Ace2 and Brg1, thereby downregulating filamentation (8), and it regulates the hypoxia-specific expression of numerous genes. Furthermore, by counteracting expression of *WOR1*, Efg1 prevents switching to the *opaque* form and favours the yeast morphology (9). The activity of the Efg1 protein is regulated by posttranslational modifications including phosphorylation by cAMP-dependent protein kinase A (PKA) in response to environmental cues (10, 11). The overall activity of Efg1 is required for biofilm formation (12, 13, 14) and virulence (2) of *C. albicans.*

In eukaryotes, the level, processing, localization and/or structure of the primary transcript determine the initial amount of the encoded protein, which is subsequently lowered by different rates of proteolytic degradation. Some of such posttranscriptional processes and their underlying mechanisms have been described in *C. albicans* to regulate levels of proteins including transcription factors (15, 16). Transcript degradation involves poly(A) tail removal by deadenylase subunits Ccr4/Pop2 (17), hydrolysis of the 5’ cap by decapping activators Dhh1/Edc3 (18), decapping enzyme Dcp1 (18) and mRNA digestion by exonuclease Xrn1/Kem1 (19, 20). RNA binding proteins Puf3 (21) and Zfs1 (22) also appear to be involved in decay of transcripts. Mutants lacking these degradative activities show defects in filamentation and/or biofilm formation, although specific targets have not yet been defined. The specific degradation of the transcript encoding Nrg1, a strong repressor of filamentation, was described to depend on an antisense transcript that originates from the locus encoding the Brg1 hyphal activator (23). The localization of transcripts also regulates filamentation of *C. albicans*, as was shown for the She3 protein that binds several transcripts involved in filamentation and transports them to the bud site of yeast cells or to the tips of hyphae (24); the Sec2 protein operating at the hyphal tip appears to specifically localize its own transcript to this location (25). It is assumed that localized translation procures directed delivery of such proteins to their sites of action. In recent years, the localization, degradation and/or translation of certain transcripts was found also to depend on promoter sequences, suggesting that already during transcription, regulatory factors for these functions may become loaded onto the emerging transcript (26-28).

The structure of the 5’ UTR of transcripts controls translation in eukaryotes. Strong evidence supports the importance of AUG context sequence on translational initiation (29, 30). Upstream open reading frames (uORFs) within the 5’ UTR can control translation of the downstream main ORF (31), as has been described for the *C. albicans GCN4* gene that regulates the amino acid starvation response, as well as filamentation and biofilm formation (32). Cap-independent translation that is initiated at internal ribosome entry sites (IRES) has been described for gene transcripts responsible for invasive growth in the yeast *S. cerevisiae* (33). In addition, 5’ UTR sequences may contain binding sites for binding proteins that facilitate localization (34) and potentially translation of transcripts. In *C. albicans*, the Dom34 protein, known for its general role in *no go* decay of mRNAs, was also shown to bind the 5’ UTR of specific transcripts encoding Pmt-type mannosyl transferases and favour their translation (35). Similarly, the Ssd1 RNA binding protein may positively affect translation of specific sets of transcripts involved in cell wall integrity and polarized growth (36, 37). Remarkably, many transcripts encoding essential regulators of cell morphology contain extensive 5’ UTR regions including *UME6* (3041 nt), *CZF1* (2071 nt), *WOR1* (2978 nt) and *EFG1* (1139 nt of long transcript) (38). The long 5’ UTRs of *UME6* and *WOR1* genes were recently shown to downregulate translation of their transcripts (39, 40), possibly by forming a tight three dimensional structure that blocks scanning by ribosomal 40S subunits. In both cases, regulated release of translational blockage may be mediated by host environmental cues that alter the 5’ UTR structure (41), e. g. in the presence of specific RNA binding proteins. Non-native, functional expression of *EFG1* has been achieved by placing the *EFG1* ORF (without the 5’ UTR sequence) downstream of the heterologous *C. albicans PCK1* and *ACT1* promoters (1, 3, 42, 43). Here we report that the extensive 5’ UTR of the major *EFG1* transcript nevertheless has a significant positive role for the functional expression of the *EFG1* ORF. A specific sequence within the 5’ UTR is required to stimulate translation of the *EFG1* transcript, to permit efficient hyphal morphogenesis.

## Results

### Deletions in the 5’ UTR of *EFG1*

In the yeast growth form (*white*), the transcript start sites for the main *EFG1* transcript are known to cluster around position −1100 relative to the ATG of the *EFG1* ORF, generating a transcript of 3.3 kb (6, 42, 44). Referring to the sequence of ATCC10231 (used here for deletion analysis), start sites lie at positions −1170, −1143 and −1112 (amended from Tebarth *et al.*, 2003), or at −1125 (−1117 in strain SC5314; Bruno *et al.*, 2010); in agreement, the start site in strain WO-1 was mapped at position −1173 (44) (Fig. 1). In the rod-like *opaque* growth form, however, low levels of a shortened *EFG1* transcript of 2.2 kb transcript occur (42), for which start sites at positions −145 and −162 were identified (44) and a start position of −74 was also observed for a minor fraction of the *EFG1* transcript in yeast-form cells (6).

**Fig. 1.**
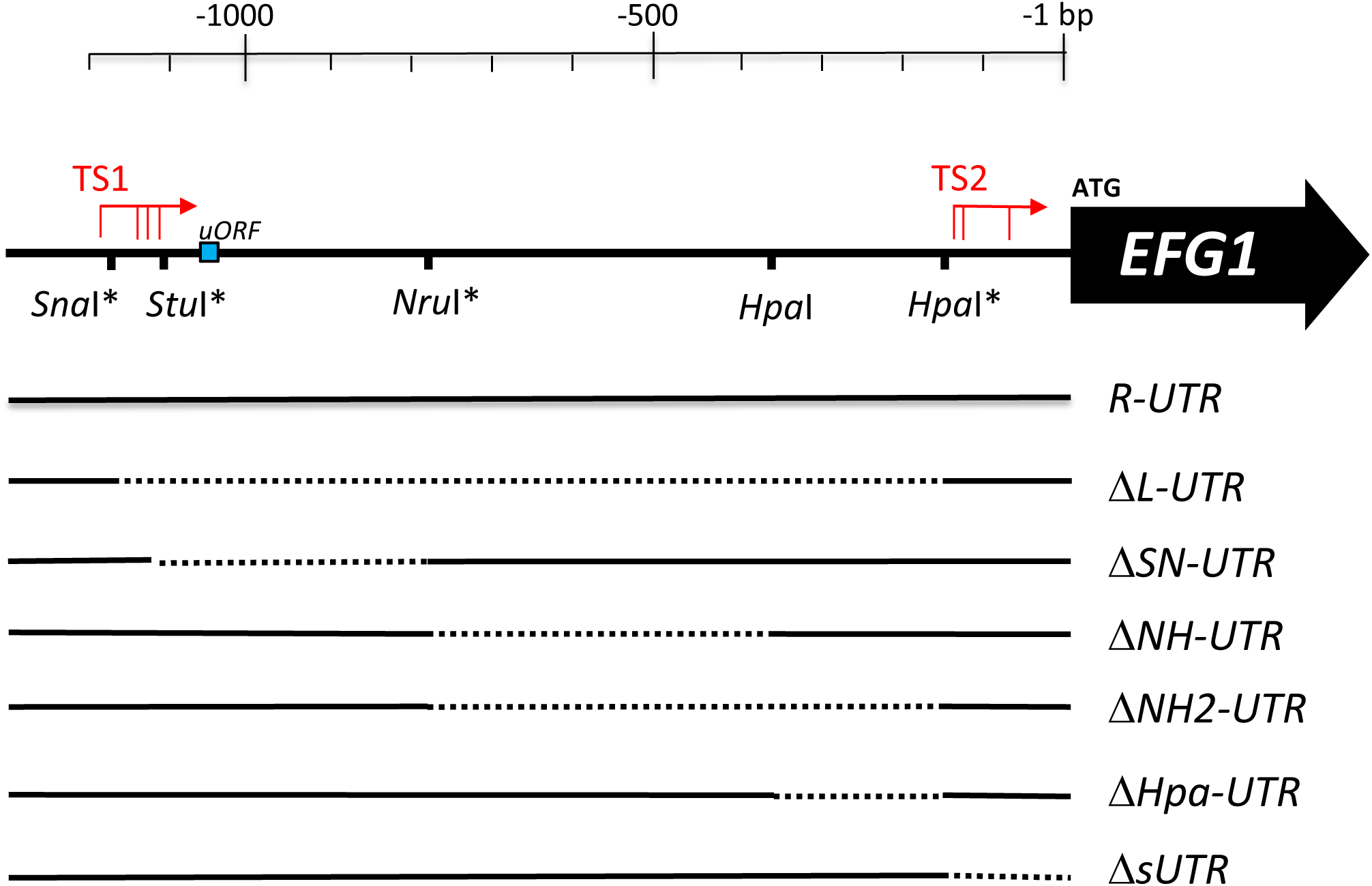
Upstream region of the *EFG1* gene. Schematic representation of the *EFG1* upstream region in strain ATCC10231 indicating start positions of the large transcript (TS1) around position −1100 and of the small transcript (TS2) around position −100. The large and small transcripts are the major transcripts observed in *white* (yeast) and *opaque* growth forms, respectively (Fig. S1). A small upstream open reading frame (uORF) encoding 4 amino acids is shown as blue box; it is missing in strain SC5314. Positions of restriction sites used to construct deletions in the 5’ UTR sequence are as indicated; sites marked by asterisks were introduced by site-specific mutagenesis. While *R-UTR* denotes the full-length 5’ UTR-*EFG1* region, the Δ series shows deleted *EFG1* alleles lacking sequences between restriction sites in the 5’ UTR (dotted lines), affecting mostly the large transcript but also the small trancript (ΔsUTR). Plasmids harbouring native and deleted forms of *EFG1* were integrated into the *EFG1* upstream region of *efg1*/*efg1* mutant HLC67.

To construct deletions in the 5’ UTR sequence, restriction enzyme sites were inserted, singly or in combination, into a plasmid-resident *EFG1* gene including 3.2 kb of its upstream sequence (allele *R-UTR)*. Sequences between restriction sites were deleted, resulting in six deleted *EFG1* alleles lacking 5 ′UTR sequences of the large transcript (Δ*L-*, Δ*SN*-, Δ*NH*-, Δ*NH2*, Δ*Hpa-UTR*) or the small transcript (Δ*s-UTR*) (Fig. 1; Fig. S1). The resulting plasmids were chromosomally integrated into the upstream region of the *EFG1* locus in strain HLC67 (2), which lacks the *EFG1* ORF (but retains its upstream sequences) on both homologous chromosomes.

### 5’ UTR sequence enhances filamentation

*C. albicans* mutants lacking the Efg1 protein are unable to form hyphae at 37 °C under all conditions, while at temperatures <35 °C, if cells are grown under hypoxia on agar surfaces, their filamentation is derepressed (4). This dual function of Efg1 as activator or repressor of morphogenesis becomes apparent during surface growth of cells under hypoxia (0.2 % O_2_) at either 25 °C or 37 °C (Fig. 2). Cells carrying at least one functional *EFG1* allele are able to filament at 37 °C but not at 25 °C, while non-functional alleles are hyperfilamentous at 25 °C, but not at 37 °C. The only exception to this pattern, as described previously (8), is mediated by the *HA-EFG1* allele, which promotes hypha formation at 37 °C but lacks repressor function at 25 °C, thus leading to filamentation under both temperatures.

**Fig. 2.**
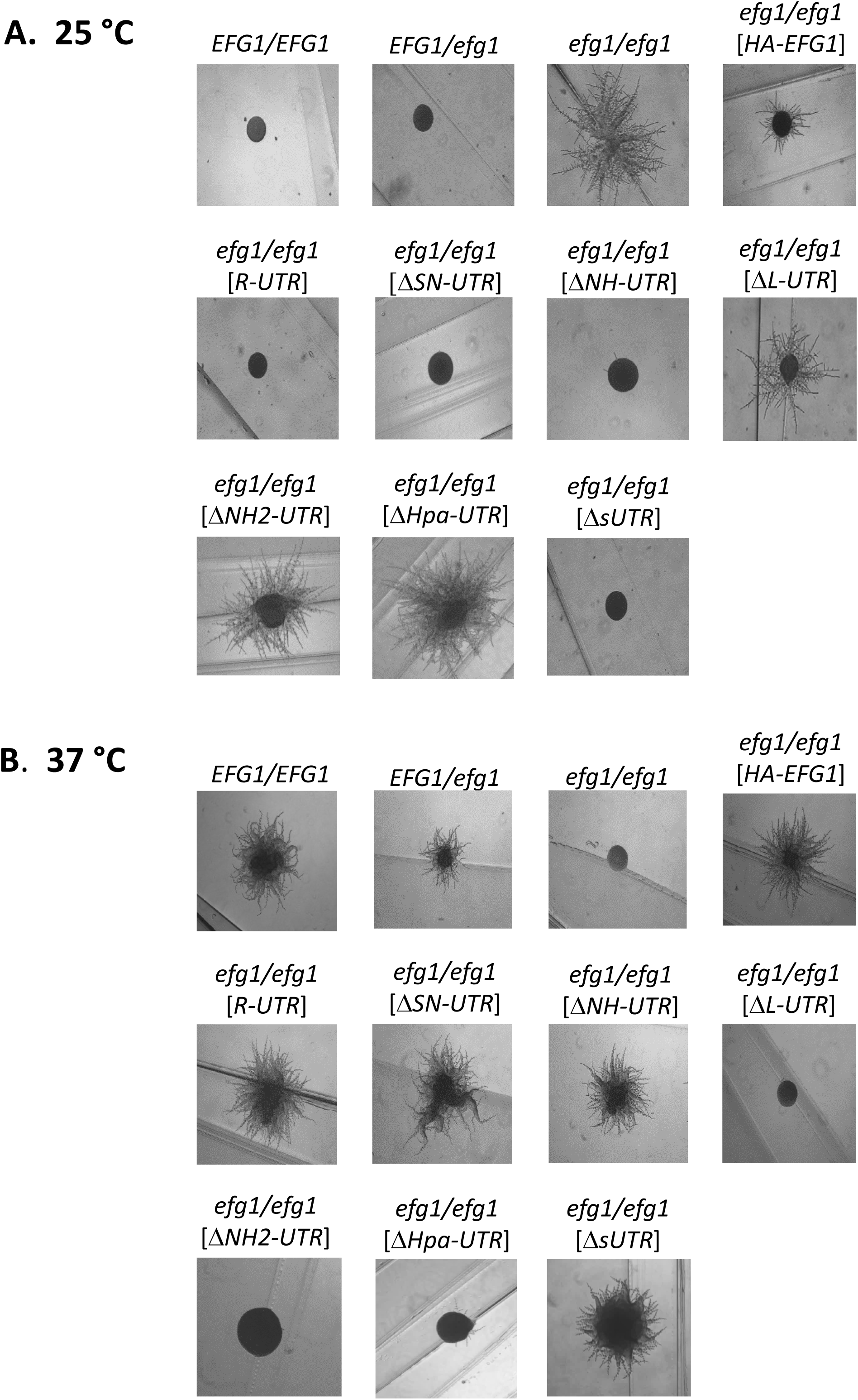
Colony phenotypes of strains expressing deletions in the 5’ UTR of *EFG1*. Strains CAF2-1 (*EFG1*/*EFG1*), BCA09 (*EFG1*/*efg1*), HLC67 (*efg1/efg1*), HLCEEFG1 (*HA*-*EFG1*/*efg1*), PDUWT (*efg1/R-UTR-EFG1*), PDUSN (*efg1/*Δ*SN-UTR-EFG1*), PDUNH (*efg1/*Δ*NH-UTR-EFG1*), PDULG (*efg1/*Δ*L-UTR-EFG1*), PDUSH (*efg1/*Δ*NH2-UTR-EFG1*), PDUHH (*efg1/*Δ*Hpa-UTR-EFG1*) and PDUsU (*efg1/*Δ*sUTR-EFG1*) were grown on Spider medium either (**A**) at 25 °C for 3 days or (**B**) at 37 °C for 2 days, under hypoxic conditions (0.2 % O_2_). The data show representative colony morphologies, which were imaged by light microscopy.

*EFG1* alleles containing either the full-length 5’UTR (*R-UTR*) or deleted alleles Δ*SN-UTR*, Δ*NH-UTR*, and Δ*sUTR* were fully active in promoting filamentation at 37 °C and repressing it at 25 °C (Fig. 2). Because deletions in these alleles encompassed a small uORF sequence, it appears that its presence is not required for hypha formation. In contrast, alleles containing Δ*L-UTR*, Δ*NH2-UTR* and Δ*Hpa-UTR* performed as non-functional *EFG1* alleles that did not stimulate filamentation at 37 °C, but allowed strong filamentation at 25 °C. The latter alleles were all lacking the 218 bp *Hpa*I fragment that was solely deleted in the Δ*Hpa-UTR* allele. To confirm these results, the function of the various alleles was also tested under normoxia using liquid induction medium containing 10 % serum at 37 °C, which demonstrated similar filamentation phenotypes as those that were observed during surface growth (Fig. 3). Thus, these results indicate that the 218 nt *Hpa*I fragment in the 5’ UTR of *EFG1* is required for production and/or activity of Efg1, promoting filamentation at 37 °C and repressing it at 25 °C. Filamentation phenotypes obtained for all tested *EFG1* alleles are summarized in Fig. 4.

**Fig. 3.**
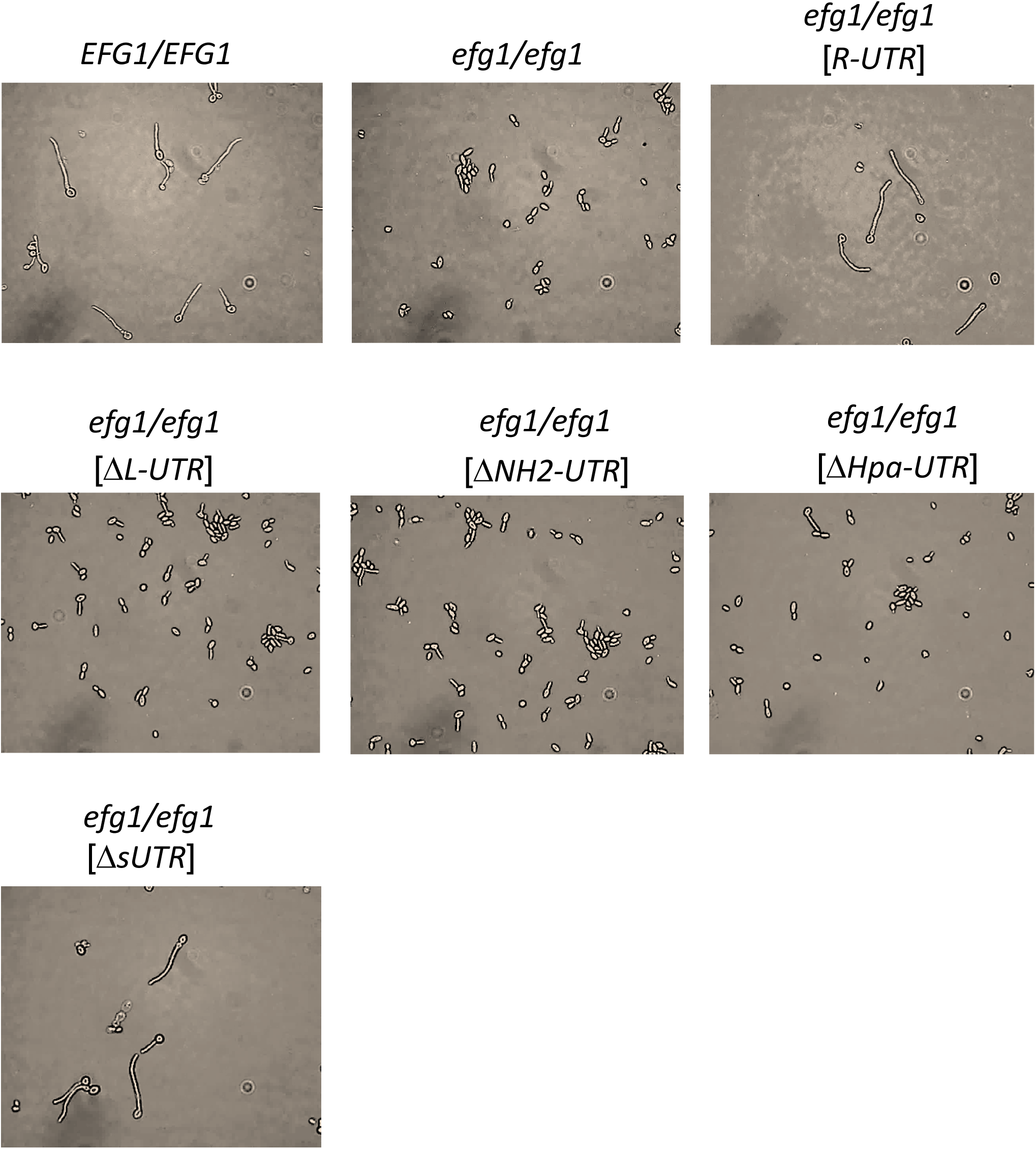
Cell morphologies of strains expressing deletions in the 5’ UTR of *EFG1* after serum induction. Strains were grown in YPD at 30 °C and diluted into pre-warmed YP medium containing 10 % horse serum at 37 °C. Cells were incubated for 30 min at 37 °C and imaged by phase contrast microscopy. Strain designations as in Fig. 2.

**Fig. 4.**
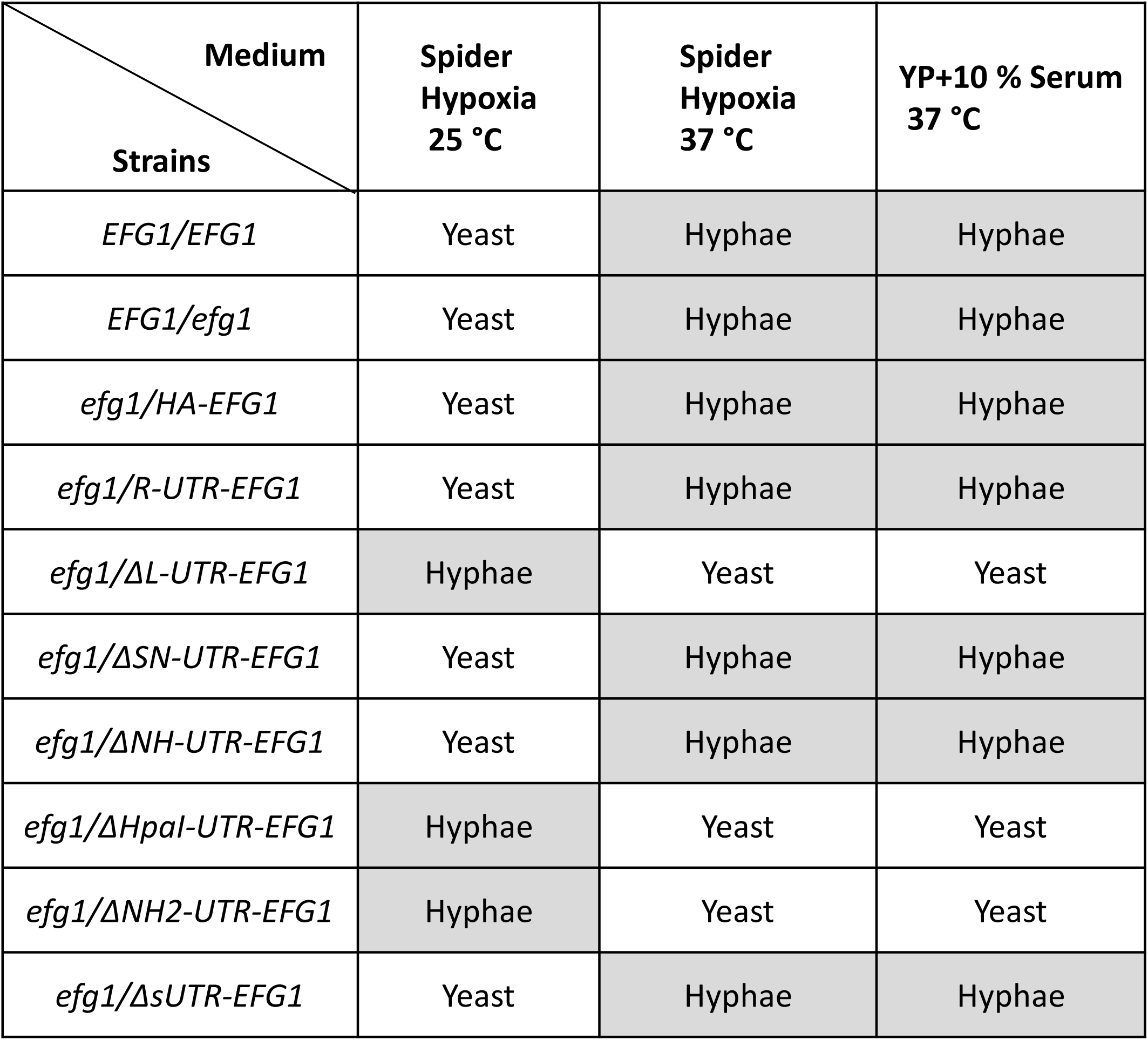
Summary of filamentation phenotypes of *C. albicans* strains carrying deletions in the 5’ UTR of *EFG1.*

### Deleted 5’ UTR alleles do not lower *EFG1* transcript levels

To clarify the reasons for the inactivity of *EFG1* alleles in cells lacking the 5’ UTR completely (Δ*L-UTR*) or partially (Δ*Hpa-UTR*), *EFG1* transcript levels were determined by qPCR. Both shortened alleles resulted in significantly elevated transcript levels, compared to wild-type cells (*EFG1/EFG1*), or to cells expressing the *R-UTR* allele containing the full-length 5’ UTR (Fig. 5 A). The observed increase was highest in cells pregrown for 12 h in YPD (t = 0), but clearly apparent also after short term growth for 2 and 4 h. It can be concluded that the low Efg1 activity of the Δ*L-UTR* or Δ*Hpa-UTR* alleles cannot be explained by lowered *EFG1* transcript levels. To verify that the respective transcripts were intact, cellular RNA was also examined by Northern blotting. As expected, wild-type cells and cells containing the *R-UTR* allele contained an *EFG1* transcript of about 3.2 kb (6, 42, 44), while the *efg1* mutant was lacking this transcript (Fig. 5 B). Remarkably, the mutated alleles encoded *EFG1* transcripts with sizes reflecting the extent of 5’ UTR deletions, i. e. the size of the transcript encoded by the Δ*Hpa-UTR* allele was only slightly reduced, while the Δ*L-UTR* transcript was shortened to about 2 kb, approximating the size that occurs in *opaque*-type cells (42, 44). These results indicate that the *EFG1* transcript encoded by the inactive, deleted 5’ *EFG1* alleles is not differentially processed or degraded.

**Fig. 5.**
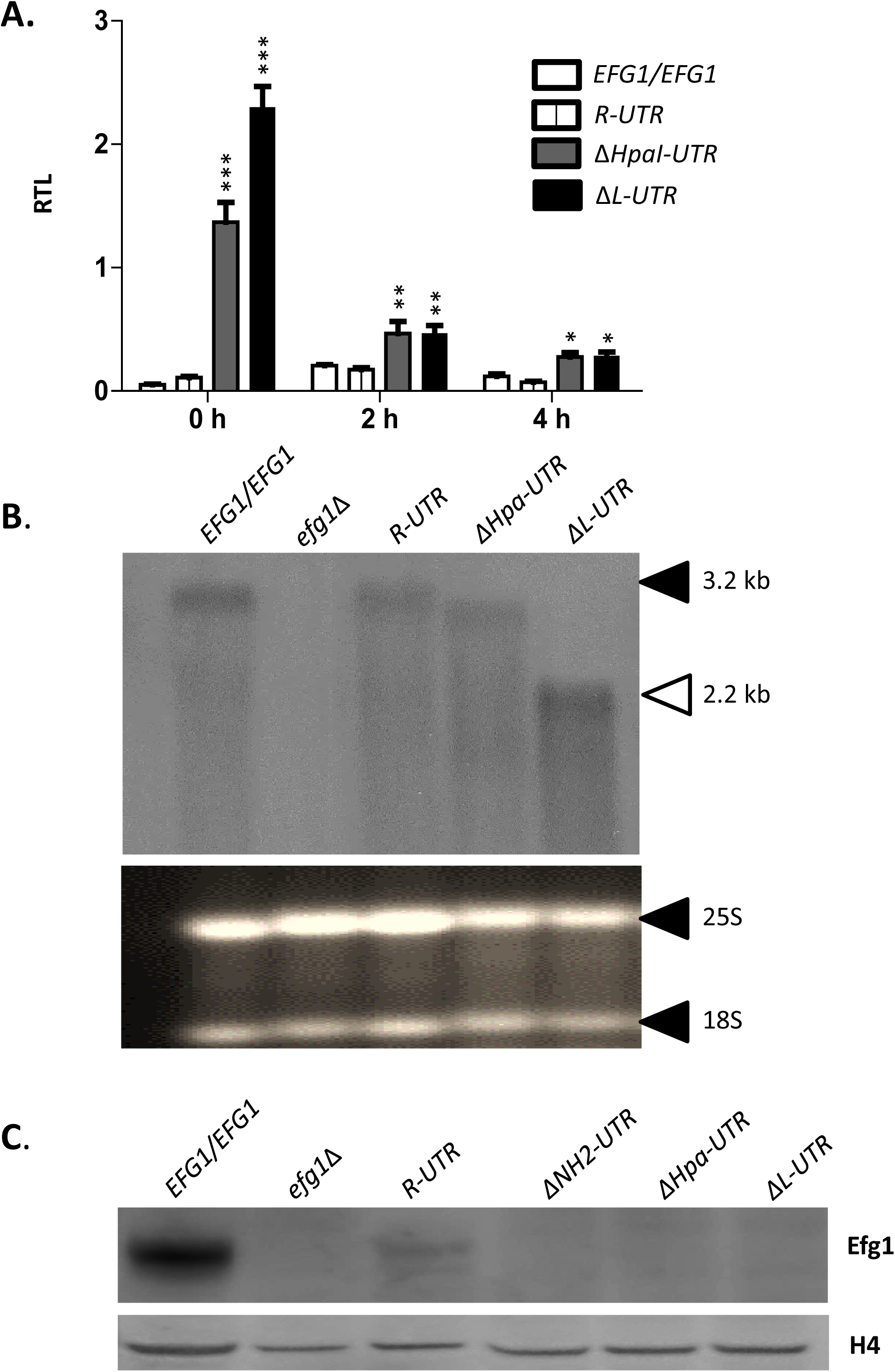
A 5’ UTR deletion increases *EFG1* transcript and decreases Efg1 protein levels. Strains CAF2-1 (*EFG1*/*EFG1*), PDUWT(*R-UTR*), PDUHH (Δ*Hpa-UTR*) and PDULG (Δ*L-UTR*) were examined for *EFG1* transcript and Efg1 protein levels. **A.** Strains were pregrown for 12 h in YPD (t = 0 h), diluted into YPD and grown 2 and 4 h at 30 °C; levels of the *EFG1* transcript were determined by qPCR, using the *ACT1* transcript as internal reference to calculate relative transcription levels (RTL). Error bars display standard error of the mean derived from biological triplicates. A two-tailed, unpaired *t* test comparing the RTL values of control CAF2-1 and other strains was used to determine the statistical relevance. *, *P* < 0.05; **, *P* < 0.01; ***, *P* < 0.001. In addition, the *EFG1* transcript in the RNA of strains grown for 6 h in YPD was examined by Northern analysis (**B**, top panel), using 8 µg RNA for loading. Note that the *EFG1* transcript size (3.2 kb) is reduced greatly only for the ΔL-UTR variant, which lacks most of the 5’ UTR region. 25S (3.4 kb) and 18S (1.8 kb) rRNA (58) stained by ethidium bromide was used as loading control (**B**, bottom panel). **C.** To determine Efg1 protein levels, strains were grown in YPD medium at 30 °C to the logarithmic phase and cell extracts derived from 1 OD_600_ equivalent of cells were separated by SDS-PAGE and analysed by immunoblotting, either using anti-Efg1 antibody or anti-histone H4 antibody for probing. Levels of histone H4 served as loading controls.

### Efg1 protein produced by deleted 5’ UTR alleles

To verify Efg1 protein levels produced by the deleted 5’ UTR alleles, cell extracts were analysed by immunoblotting, using an anti-Efg1 antiserum described previously (7, 45). The Efg1 protein was detected strongly in wild-type cells (carrying two *EFG1* alleles) and also, with reduced intensity in cells carrying a single *R-UTR* allele containing the full-length 5’ UTR (Fig. 5 C). In contrast, no Efg1 protein was observed in cells expressing the trucated 5’ UTR versions Δ*L-UTR*, Δ*Hpa-UTR* or Δ*NH2-UTR*, which are functionally inactive. It can be concluded that the latter alleles do not produce significant amounts of Efg1 protein, in spite of expressing high *EFG1* transcript levels.

### Truncation of the 5’ UTR deletion reduces translation of *EFG1*

The above results had suggested that the 5’ UTR of the *EFG1* transcript contains a 218 nt sequence corresponding to the small *Hpa*I fragment of the *EFG1* upstream region, which is required for efficient translation of Efg1. To test this hypothesis polysome analyses were carried out using cellular lysates of strains expressing *EFG1* alleles containing the full-length 5’ UTR (*R-UTR*) or the partially deleted variant (Δ*Hpa-UTR*). As expected, profiles obtained by sucrose gradient centrifugation were similar in both strains, showing a pre-polysomal fraction (containing 40S, 60S and 80S rRNA) and several polysomal peaks (Fig. 6 A). Transcript levels of *EFG1* and the *ACT1* housekeeping gene in the pre-polysomal and polysomal fractions were examined by qPCR, using a spiked-in control RNA as a reference. The results demonstrate that the *EFG1* transcript containing the full-length 5’ UTR is significantly enriched in the polysomal fraction compared to the pre-polysomal fraction (Fig. 6 B), while in cells expressing the Δ*Hpa-UTR* allele the *EFG1* transcript occurred in similar amounts in pre- and polysomal fractions. In contrast, the *ACT1* transcript used as a control was increased in the polysomal fraction and occurred in similar amounts in both types of cells (slightly increased in cells with the Δ*Hpa-UTR* allele). The results indicate that a specific deletion within the 5’ UTR of the *EFG1* transcript impairs its translation.

**Fig. 6.**
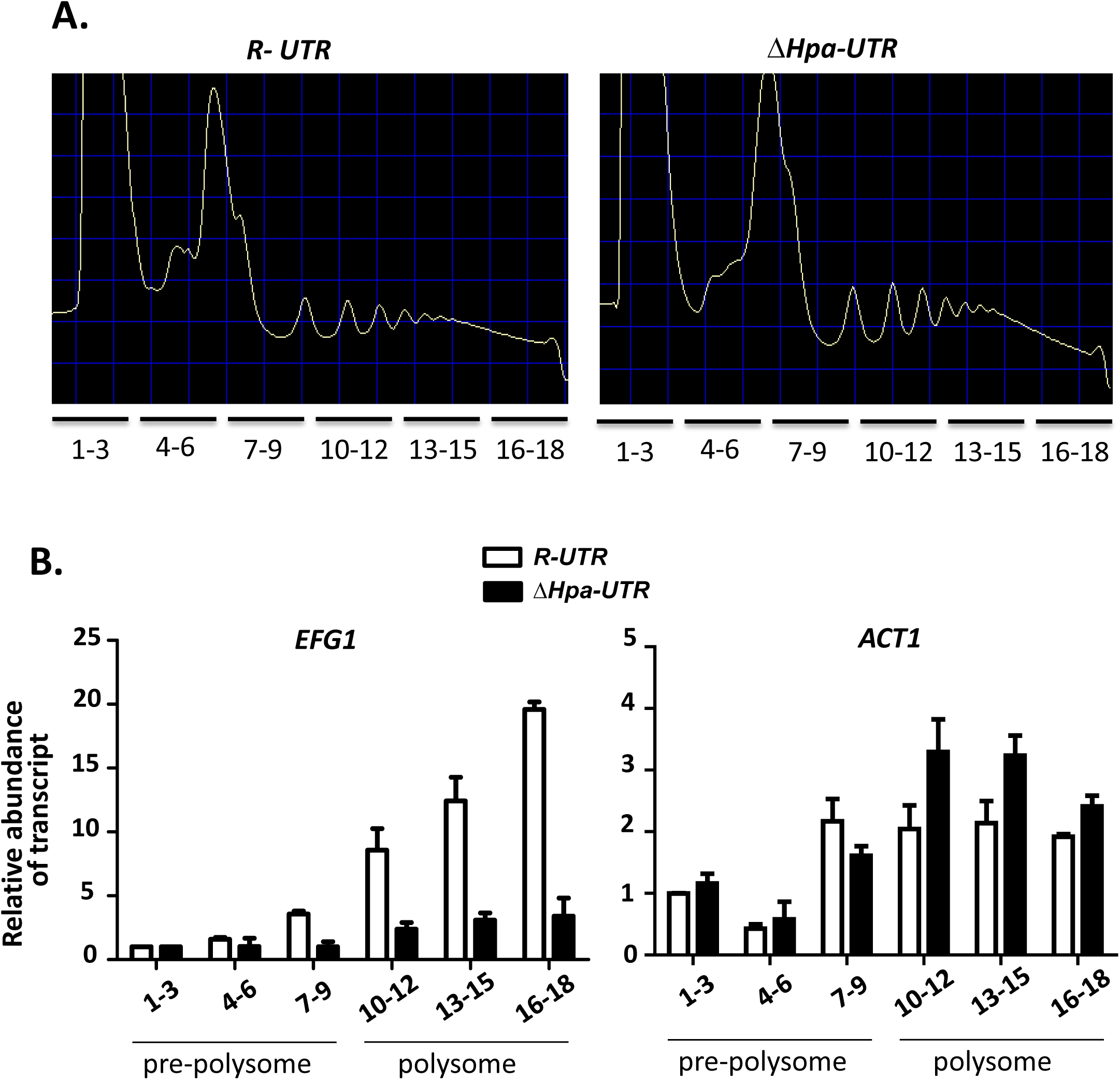
Transcript fractionation on polysome gradients. **A.** Strains PDUWT (*R-UTR*) and PDUHH (*ΔHpa-UTR*), pregrown in YPD medium at 30 °C, were transferred to YP medium containing 10 % horse serum at 37 °C and incubated for 15 min. Cellular extracts of strains were centrifuged in a 10-50 % sucrose gradient, which was subsequently fractionated. Nucleic acids in gradient fractions were detected by absorbance (A_260_). Note that pre-polysome fractions 1-9 contain 40S, 60S and 80S ribosomal RNA. **B.** *EFG1* and *ACT1* transcripts in gradient fractions were detected by qPCR after adding a known amount of an *in vitro* generated transcript of the *CaCBGluc* gene as calibrator. Data shown represent values that are normalized to *EFG1* or *ACT1* mRNA abundance in fraction 1. Each bar represents the normalized mean *EFG1* or *ACT1* transcript level of two independent experiments with three technical replicates and includes the standard error of the mean.

### ORF-independent function of the 5’ UTR sequence

The observed positive effect of the 5’ UTR of the *EFG1* transcript on its translation could either operate independently, or dependent on its native context upstream of the *EFG1* ORF. This possibility was examined by replacing the *EFG1* ORF in control strain PDUWT (*efg1/R-UTR-EFG1*) by the heterologous *CaCBGluc* sequence that encodes click beetle luciferase (46); thereafter, the resulting strain EFG1GN contained the allele *EFG1p*-*R-UTR-CaCBGluc*. Likewise the *EFG1* ORF was replaced in strain PDUHH (*efg1*/Δ*Hpa-UTR-EFG1*), resulting in strain DUTRinEFG1GN containing allele *EFG1p*-Δ*Hpa-UTR-CaCBGluc*. As controls, the *CaCBGluc* gene was also used to replace one allele of the *ACT1* ORF in both PDUWT and PDUHH, generating strains ACT1GN and DUTRinACT1GN that both carry the *ACT1p*-*CaCBGluc* fusion. *CaCBGluc* transcript levels driven by the *ACT1* promoter were similar in strains ACT1GN and DUTRinACT1GN, as expected (Fig. 7 A); correspondingly, luciferase activities were nearly identical (Fig. 7 B). Under control of the *EFG1* promoter that was joined to the intact 5’ UTR (*R-UTR*), the *CaCBGluc* transcript level was about 5 fold higher as compared to its junction to the deleted 5’ UTR sequence (allele Δ*Hpa-UTR*), suggesting that truncation of the 5’ UTR lowers transcript stability. It should be considered here that negative autoregulation known for the *EFG1* gene (Fig. 5) (6, 7) cannot occur for the described *CaCBGluc* fusions. Remarkably, however, in spite of considerable *CaCBGluc* transcript levels, luciferase activity was essentially lost in strain DUTRinEFG1GN. The complete loss of luciferase activity was surprising, considering that the *CaCBGluc* transcript level in this strain was even higher than in control strain DUTRinACT1GN (*CaCBGluc* transcribed by the *ACT1* promoter), which generated abundant luciferase activity. The results support the importance of the 5’ UTR *EFG1* sequence for the functional expression of the downstream ORF, which need not be the native *EFG1* ORF.

**Fig. 7.**
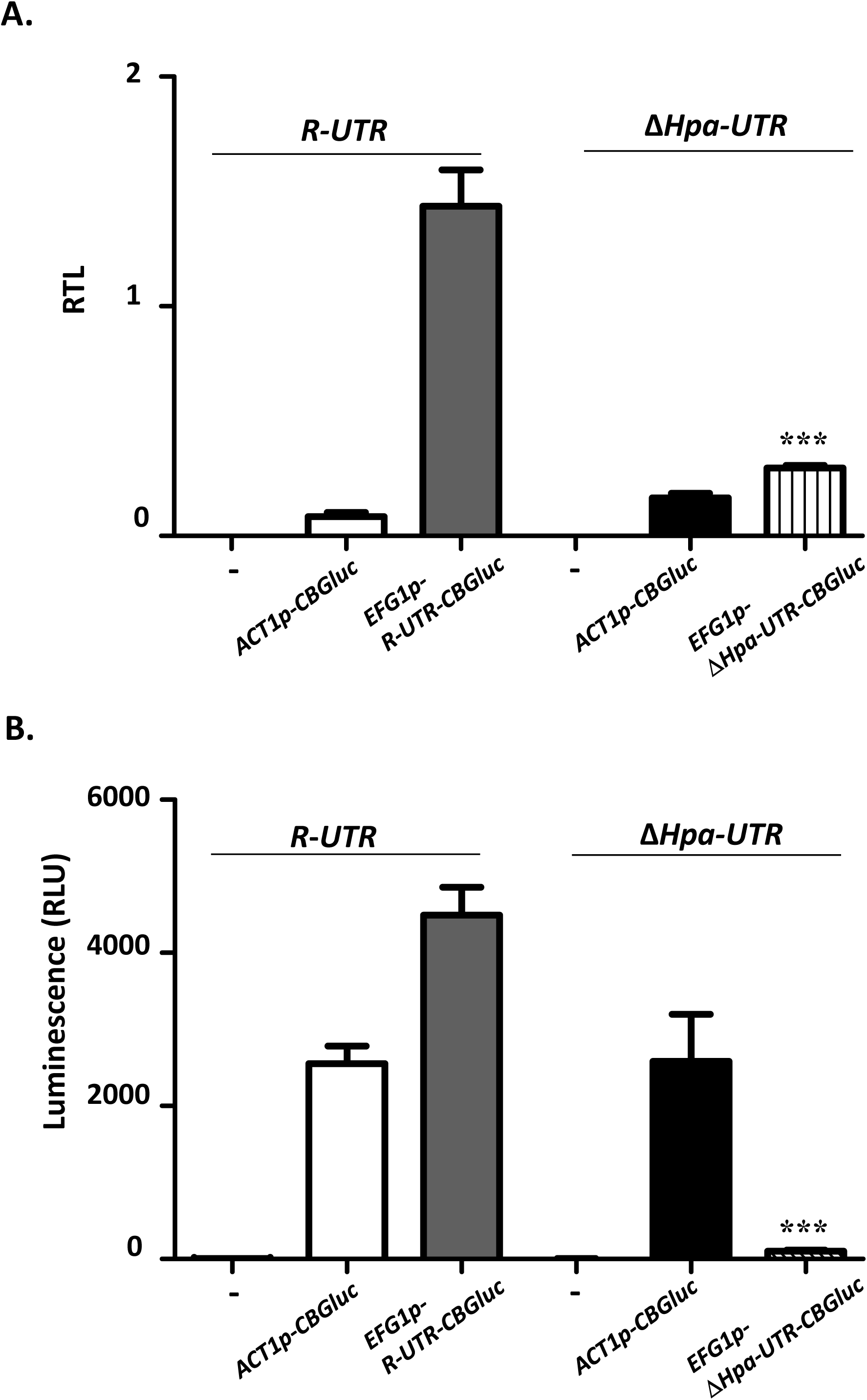
Deletions in 5’ UTR affect the translation of *CaCBGluc*. Strains either containing the intact *R-UTR* allele (strain PDUWT) or the Δ*Hpa-UTR* allele (strain PDUHH) of *EFG1* were modified further by replacing either the *ACT1* or *EFG1* ORF by the *CaCBGluc* ORF. Resulting strains include ACT1GN (*efg1/EFG1p-R-UTR-EFG1, ACT1/ACT1p*-*CaCBGluc*), EFG1N (*efg1/EFG1p-R-UTR-CaCBluc*), DUTRinACT1GN (*efg1*/*EFG1p-*Δ*Hpa-UTR-EFG1, ACT1/ACT1p-CaCBGluc*) and DUTRinEFG1GN (*efg1*/*EFG1p-*Δ*Hpa-UTR-CaCBGluc*). Strain PDUWT (*efg1/EFG1p-R-UTR-EFG1*) was used as control (−). Strains were grown in YPD medium at 30 °C to the exponential phase (OD_600_= 0.5) and used for transcript and luciferase activity levels. **A.** *CBGluc* transcript level. Relative transcript levels (RTL) of *CaCBGluc* were determined by qPCR, using the *ACT1* transcript for normalization. Error bars display standard errors of the mean derived from three biological and three technical replicates. **B.** Luciferase activity. Luminescence originating from 100 µl of cells was assayed after addition of 100 µl of Beetleglow reagent. Statistical significance was determined by comparing the EFG1GN and UTRinACT1GN strains, with a two-tailed, unpaired *t* test based on two biological and three technical replicates. *, *P* < 0.05; **, *P* < 0.01; ***, *P* < 0.001.

## Discussion

The dual activity of Efg1 as an activator and repressor of transcription requires proper timing and targeting of its activity. Although Efg1 is required to initiate hypha formation under normoxia (1,2), its prolonged activity interferes with orderly filamentation (6, 7). Under some hypoxic conditions, Efg1 is not an activator, but an efficient repressor of hypha formation (3, 4). Efg1 induces genes specific for the yeast (*white*) growth form, but by repressing *WOR1*, it prevents the rod (*opaque*) growth form (9). In metabolism, Efg1 induces genes involved in glycolysis, but it also represses genes in oxidative metabolism (3). Furthermore, Efg1 induces and represses hypoxia-specific genes, and it prevents inappropriate hypoxic regulation of genes not normally regulated by oxygen (4). Efg1 activity is hitherto known to be regulated on posttranslational and transcriptional levels. Posttranslational modes of regulation include Efg1 phosphorylation by PKA isoforms (10, 11), which may occur directly at target genes (47) or physical association with regulatory factors like Flo8 and Czf1 (45). Transcriptional repression of *EFG1* expression is mediated by Sin3 (6), Wor1 (9) and also by Efg1 itself (6,7), causing negative autoregulation that prevents an overshoot of Efg1 activity. *EFG1* activation is mediated in an environment-dependent manner by Brg1, Bcr1 of Ace2 (8). Here, we report a novel mechanism regulating Efg1 biosynthesis on the translational level.

We present evidence that a 218 nt sequence of the 5’ UTR of its major transcript is required for Efg1 protein production. Because of negative autoregulation of *EFG1* (6, 7), transcript levels of the 218 nt deletion variant were even increased, but still did not yield significant amounts of Efg1 protein. In wild-type cells, the major *EFG1* transcript was distributed mostly to polysomes, while the deleted transcript was equally distributed to monosomes and polysomes, suggesting that the 218 nt sequence activates Efg1 translation. This positive effect was even observed, if the *EFG1* ORF was replaced by the ORF of a heterologous reporter gene, indicating that the activating function of the 5’ ORF does not depend on its native 3′ context. As expected from these results, the absence of this regulatory sequence in the short 2.2 kb transcript of the *opaque* form (or in the minor 2.2 kb transcript of the *white* yeast form) (6, 42, 44) is expected to reduce the production of Efg1 protein. This mechanism contributes to lowering Efg1 activity in *opaque* cells, which is already reduced on the transcriptional level (9, 42, 44), to prevent backward switching to the *white* (yeast) form. Clearly, low translation of the *EFG1* transcript in *opaque* cells (40), is not caused by an inhibitory effect of the 5’ UTR, as has been suggested (40), but it is due to the lack of the 218 nt sequence in the short *opaque* transcript (2.2 kb). The positive translational function of the 5’ UTR in the *EFG1* major transcript differs from other recently reported 5’ UTR regions in transcripts of two different *C. albicans* genes. In contrast to *EFG1*, 5’ UTR sequences of both *UME6* and *WOR1* transcripts were found to negatively influence translation of the respective proteins (39, 40). Furthermore, both *UME6* and *WOR1* are positively autoregulated (39, 40, 48), while *EFG1* is negatively autoregulated. The different modes of autoregulation nevertheless lead to increased promoter activities and transcript levels of all three genes lacking the 5’ UTR (or relevant parts thereof): in the case of *UME6/WOR1* this result is caused by relief of translational inhibition (increased protein levels stimulate promoter activity), while for *EFG1* this occurs because Efg1 production is reduced, which derepresses *EFG1* promoter activity.

The molecular mechanism, by which the 5’ UTR sequences of *EFG1* or *UME6/WOR1* transcripts regulate translation, is not known and needs experimental verification. The 218 nt sequence of the *EFG1* 5’ UTR is predicted to form a hairpin (Fig. 8), which possibly could help to generate a mRNA structure that is favourable for translational initiation. This potential structure could also be the target of RNA binding proteins that stimulate translation. Recently, the *C. albicans* Dom34 protein, a predicted component of the *no go* transcript degradation pathway, was found to bind to the 5’ UTR of transcripts encoding protein *O*-mannosyltransferases and to promote their translation (35). Binding proteins could also have an inhibitory function, such as the Rim4 protein in the yeast *S. cerevisiae* that binds to the 5’ UTR of the *CLB3* transcript to inhibit its translation (49). Likewise, the Ssd1 protein represses translation of genes involved in cell growth and morphogenesis by binding to the 5’ UTR of target transcripts (36). In mammalian cells, glucose-induced translation of insulin requires proteins binding to the 5’ UTR of the encoding transcript (50). On the other hand, the 5’ UTR structure of several human gene transcripts are known to mediate translational control that is essential to prevent several serious diseases (51). The function of 5’ UTR binding proteins is possibly related to the regulation of ribosomal assembly at the AUG initiation codon. Interestingly, the recruitment of regulatory factors to transcripts may not only depend on 5’ UTR or other transcript sequences, since promoters also can provide regulatory proteins that control the degradation, localization and translation of transcripts (26, 27). It has been suggested that such proteins may be loaded onto the mRNA near its 5’ end early in transcription (28). Such a mechanism could also be operative for the *EFG1* 5’ UTR, because its positive effect was only detected in the context with its native upstream promoter sequences, but not with heterologous *PCK1* and *ACT1* promoters, which were able to drive functional expression of the *EFG1* ORF lacking the 5’ UTR (1, 3, 42, 43). Although the functional interplay of promoter and 5’ UTR sequences remains to be established, it is possible that *EFG1* promoter sequences support the action of the 5’ UTR in translation, e. g. by transcript loading with positively-acting translation factors. Several other mechanisms explaining the regulatory function of the 5’ UTR sequence in the major *EFG1* transcript are possible. Internal ribosome entry sites (IRESs) have not only been described for viral transcripts or genomes, but also for translation of yeast genes involved in responses to starvation, which require IRES sites within transcripts (33). uORF sequences can occupy 5’ UTRs and contribute to regulation of eukaryotic translation (31). In *C. albicans*, for example, an uORF regulates translation of the *GCN4* transcript (32). We identified a short uORF with an AUG start codon in the 5’ UTR of *EFG1* in the *C. albicans* strain ATCC2013. However, this uORF does not appear to be relevant, since it does not occur in the *EFG1* 5’ UTR of strain SC5314 and its deletion did not influence functional expression of *EFG1* in strain ATCC2013. However, it should be considered that in the yeast *S. cerevisiae* translational initiation has been observed also at non-AUG codons, especially at UUG and GUG (52) and the use of GUG for translational initiation in *C. albicans* has already been reported (53). Interestingly, assuming that UUG can be used for translational initiation in *C. albicans*, two uORFs placed side-by-side are predicted within the 218 nt regulatory sequence of the *EFG1* transcript (Fig. S1). These uORFs could potentially encode peptides of 53 and 29 amino acids, respectively. In general, however, uORFs are known to negatively influence the translation of ORF sequences that are situated immediately downstream, rather than acting positively as in the case of the *EFG1* ORF (31). Since all identified uORFs also terminate in the 5’ UTR of *EFG1*, a potential translational read-through generating an extended Efg1 protein, as has been observed for Myc (54), can be excluded. Whatever the underlying mechanism of regulation by 5’ UTR sequences maybe, it may be relevant for a significant number of virulence-related *C. albicans* genes that carry extensive 5’ UTR regions. It can also be speculated that such processes may become new targets for antifungal compounds.

**Fig. 8.**
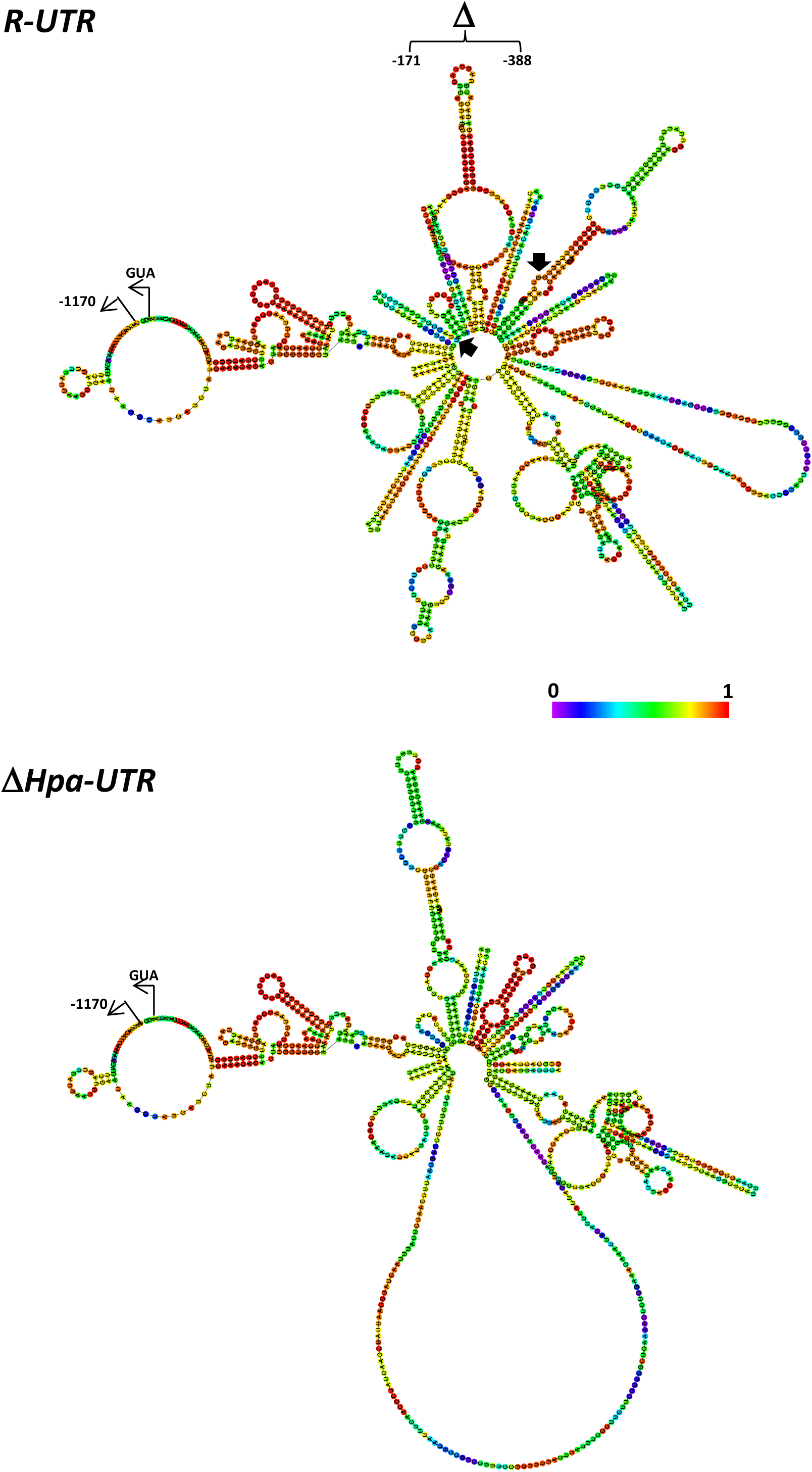
Secondary structure of *EFG1* 5’ UTR. Predicted folding structure of (**A**) full length 5’ UTR (*R-UTR*) and (**B**) deleted 5’UTR (Δ*Hpa-UTR*) of *EFG1.* The RNAfold program (http://rna.tbi.univie.ac.at/cgi-bin/RNAWebSuite/RNAfold.cgi) was used for prediction and results were depicted as a centroid structure drawing base-pair probabilities. The colour code indicates probabilities of base-pairing or single-strandedness in predicted paired and unpaired regions, respectively. Black arrows frame the region deleted in the Δ*Hpa-UTR* structure (Fig. S1), which lacks a strong hairpin between positions −327 to −229. Note that strong pairing is predicted between the 5’ end of the UTR (−1118 to −1100) and sequences preceding the AUG translational initiation codon (−112 to −25). This structure is still present in the deleted UTR.

## Material and Methods

### Strains and media

Strains used in this study are listed in Table S1. Strains were grown in liquid YP (1 % yeast extract, 1 % peptone) with either 2 % glucose (YPD) or 10 % horse serum, to induce filamentation. To induce filamentation on agar, strains were grown on Spider medium (0.3 % beef extract, 0.5 % peptone, 0.2 % K_2_HPO_4_, 1 % mannitol and 2 % agar, pH 7.2). An Invivo200 hypoxia chamber (Ruskinn) was used for hypoxic growth (0.2 % O_2_).

### Construction of strains containing deletions in the 5’ UTR of *EFG1*

Expression vector pTD38-HA (45) was modified to remove sequences encoding the N-terminal HA-tag, which has been shown to block filamentation phenotypes of Efg1 (8). For this purpose, an *Afl*II restriction site was introduced by site specific mutagenesis (QuikChange kit, Agilent) using primers MAflIIFor/rev (Table S2) downstream of the HA tag sequence, between positions −7 to −2 bp upstream of the *EFG1* ORF (sequence 5’-ACCC<u>CTTAAG</u>A ATG). The resulting plasmid pPD21HA-AB was cut with *Pac*I and *Afl*II to remove all upstream sequences, which were replaced by a fragment lacking HA sequences generated by PCR using primers 5UTREfgSphIFor/5UTREfgAflIIrev using pTD38-HA as template. The resulting plasmid pPD21-AB contains 3.2 kb of upstream sequences (comprising 2 kb of promoter and 1.2 kb of 5’ UTR sequences) upstream of the *EFG1* ORF. To delete sequences within the 5’ UTR, novel restriction sites were inserted singly or in combination by site-specific mutagenesis at position −1167 (*Sna*BI), −1112 (*Stu*I), −787 (*Nru*I), and −167 (*Hpa*I), using primers listed in Table S2 (Fig. S1). Plasmids were digested pairwise using *SnaB*I/*Hpa*I, *Stu*I/*Nru*I, *Nru*I/*Hpa*I (native *Hpa*I site at −391), *Nru*I/*Hpa*I (−167), *Hpa*I (−391)/*Hpa*I (−167) enzymes and re-ligated, to generate plasmids pΔL-UTR, pΔSN-UTR, pΔNH-UTR, pΔNH2-UTR and pΔHpa-UTR. Furthermore, the sequence between *Hpa*I (−167) and position −6 was deleted using primer mutagenesis, to construct plasmid pΔsUTR. Plasmids were linearized with *Pac*I (1.9 kb upstream of *EFG1* ORF) and transformed into strain HLC67 (*efg1* mutant lacking the *EFG1* ORF). The correct integration of plasmid in *EFG1* locus was confirmed by colony PCR using primers ColoEfg1For/ColoEfg1Rev.

### Construction of strains producing click beetle luciferase

To construct a plasmid carrying a green click beetle luciferase gene with a *sat1* nourseothricin selection maker gene, the plasmid pGEM-HIS1-CBG (46) was restricted with *Bam*HI and *Msc*I (New England Biolabs) to cut out and replace the *HIS1* gene sequence. The sequence for the *sat1* marker was optained from the donor plasmid PFA-SAT1 (55) using the two restriction enzymes *Pvu*II and *Bam*HI. The obtained *sat1* sequence was then integrated into the pGEM plasmid directly downstream of the *CBG* gene via ligation to obtain the plasmid pGEM-SAT1-CBG, which was used as the *CBGluc-sat1* reporter cassette template. Reporter cassettes were amplified via PCR with the primer pairs inACT1-CBG-Fw / inACT1-SAT1-Bw and inEFG1-CBG-Fw / inEFG1-SAT1-Bw (Table S2). These primers carry 60 bp homology to the gene of interest, *ACT1* and *EFG1*, respectively. The DNA fragments were transformed into the parental strains PDUWT (*efg1/R-UTR-EFG1*) and PDUHH (*efg1*/ΔHpa-UTR-*EFG1*). Homologous integration of the luciferase-*sat1*-reporter cassette sequence occurred downstream of the respective start codon of *ACT1* or *EFG1* genes, resulting in 2 reporter strains each for PDUWT (ATC1GN, EFG1GN) and PDUHH (DUTRinACT1GN and DUTRinEFG1GN). Mutants were selected for positive luminescence signals and correct integration was checked via colony PCR using the primer pairs ACT1 col Fw / CBG col Bw (*ACT1*) and EFG1 col Fw / CBG col Bw (*EFG1*). Mutants both positive for colony PCR and luminescence were used for further experiments.

### Blotting procedures

For Northern blottings, the strains were grown at 30 °C to the logarithmic phase, total RNA was isolated and 8 µg of RNA were separated on agarose gels containing 1.2 % formaldehyde. Following transfer to nylon membranes (Roche), blots were hybridized with ^32^P-labelled probes for *EFG1* using primers ProFor and ProRev. For signal detection the washed membranes were exposed to phosphofilm (Fujifilm) for 30 to 60 min and scanned by the Phosphor Imager FLA 5000 (Fujifilm).

For immunoblottings, YPD precultures grown over night at 30 °C in YPD medium were used to inoculate 30 ml of YPD medium. Strains were grown to OD_600_ = 0.1, harvested by centrifugation, frozen at −70°C for 1 h, and then thawed by addition of 500 ml of CAPSO buffer (20 mM CAPSO/pH 9,5, 1 M NaCl, 1 mM EDTA, 20 mM imidazole, 0,1% Triton X-100) containing protease inhibitor (Cocktail Complete, Mini, EDTA-free/Roche). Cell extracts were prepared as described in Desai *et al*., 2015). 80 μg of the crude cell extract was separated by SDS-PAGE (10 % polyacrylamide) and analysed by immunoblotting using anti-Efg1 antiserum (1: 5,000) (8) or anti-histone H4 (Abcam; 1: 5,000) to detect histone H4 as loading control. Anti-rabbit-IgG-HRP conjugate (1:10,000) was used as secondary antibody in all blottings. Signals generated by the chemiluminescent substrate (SuperSignal West Dura; Pierce) were detected by a LAS-4000 mini imager (Fujifilm) and evaluated by the Multi Gauge Software (Fujifilm).

### Polysome profiling

*C. albicans* strains PDUWT and PDUHH were grown exponentially in YPD media to OD_600_ 0.4 – 0.6. For preparation of samples derived from cells following hyphal induction, exponentially growing cells were washed with 1 X PBS and resuspended in YP medium containing 10 % horse serum (pre-warmed at 37 °C) and incubated at 37 °C for 15 min. Preparation of cells for polysome gradients was performed as described previously (35, 56), with some modifications. A portion of the culture (80 ml) was recovered and chilled for 5 min on ice in the presence of 0.1 mg/ml cycloheximide (CHX). Cells were harvested by centrifugation at 6000 *x g* for 4 min at 4 °C and resuspended in lysis buffer (20 mM Tris-HCl/pH8, 140 mM KCl, 5 mM MgCl_2_, 0.5 mM dithiothreitol, 1 % Triton X-100, 0.1 mg/ml cycloheximide, and 0.5 mg/ml heparin). After washing, cells were resuspended in 700 μl of lysis buffer, 300 µl glass beads were added and cells were disrupted by shaking on a Vortex Genie2 (setting 8) using 6 cycles for 40 sec. Between cycles, cells were placed on ice for 5 min. Lysates were cleared by centrifuging twice for 5 min, first at 5,000 rpm, and then the supernatant was recovered and was centrifuged at 8,000 rpm. Finally, glycerol was added to the supernatant at a final concentration of 5 %, before storing extracts at −70 °C. Samples of 10–20 A_260_ units were loaded onto 10 - 50 % sucrose gradients and were separated by ultracentrifugation for 2 h and 40 min at 35,000 rpm in a Beckman SW41 rotor at 4 °C. Then, gradients were fractionated using isotonic pumping of 60 % sucrose from the bottom, followed by recording of the polysomal profiles by online UV detection at 260 nm (Density Gradient Fractionation System, Teledyne Isco, Lincoln, NE). To analyse the RNA of the polysomal fractions, RNA from 200 μl of each fraction was extracted using GeneJet RNA extraction kit (STREK, Biotools).To each sample, 500 ng of *in vitro* transcribed RNA (HiScribe™ T7 High Yield RNA Synthesis Kit, NEB) was added and used as spiked-in mRNA for normalization of the transcripts. After reverse transcription of the purified RNA (Maxima First Strand cDNA synthesis kit, Thermo Scientific), quantitative PCR (RT-qPCR) was performed using gene specific primer pairs to quantify mRNAs of *EFG1* and *ACT1*. For each fraction two biological replicates with three technical replicates were assayed on a Mx3000P Light Cycler (Stratagene), with 10 μl of cDNA, 4 μl EvaGreen qPCR-mix II (Bio-Budget) and 3 μl each of forward and reverse oligonucleotide primers (400 pmol/μl) in each reaction. The polymerase was activated at 95 °C for 10 min, annealing was performed at 60 °C for 15 sec, extension at 72 °C for 30 sec and the denaturation step was performed at 95 °C for 30 sec, using a total of 50 cycles.

### qPCR

cDNA for qRT-PCR analysis was prepared from 2 μg of total RNA treated with DNAse I (Thermo Fischer) using the Maxima First Strand cDNA Synthesis Kit (Thermo Fischer). Real-time PCR was performed in triplicate in 96-well plates using the EvaGreen dye (Bio-Budget). Primers used for qRT-PCR analysis are described in Table S2. Real-time PCR was performed using the following cycling conditions: Step 1: 95 °C for 15 minutes, Step 2: 95 °C for 15 seconds, Step 3: annealing temperature 60 °C for 20 sec, Step 4 elongation : 72 °C for 20 sec, Step 5: repeat steps 2-4 for 39 times, Step 7: melting curve 50 °C – 95 °C every 0.4 °C, hold 1 sec and read plate. Expression levels of each gene were normalized to levels of an internal *ACT1* control using the Pfaffl method (57).

### Luciferase assay

To measure CB luciferase activity in yeast cells, overnight cultures of PDUWT, PDUHH, ACT1GN, EFG1N, UTRinACT1GN and UTRinEFG1GN were diluted to OD _600_= 1.0 in PBS buffer (140 mM NaCl; 3 mM KCl; 8 mM Na _2_HPO _4_; 1.8 mM KH _2_PO _4_/pH 7.4) and incubated at 30 °C for 60 min at 180 rpm. 1 ml were transferred into fresh YPD medium and grown for 6 h at 30 °C. All samples were set to an OD _600_ = 0.3 in 1 ml YPD and quickly frozen in liquid nitrogen. After thawing, 100 µl of the samples were transferred into a 96-well microtiter plate and 100 µl Beetleglow (46) was added to start the reaction. Measurements were made in a infinite M200 PRO plate reader (Tecan) with the following settings. Plates were shaken for 10 s at 140 rpm and relative luminescence units (RLU) were measured for 1 s per well at 30 °C. Each plate was measured 3 times and the maximal luminescence values (L _max_) were reported.

## Acknowledgements

This work was supported by the Infect-ERA JTC2 project FunComPath (http://www.funcompath.eu/) to J.F.E. P.A. was funded from Mineco and FEDER funds (BFU2016-77728-C3-3-P). The authors declare no conflict of interest. This paper is dedicated to the memory of André Goffeau.

## Supplemental Material

**Fig. S1. Upstream sequence of the *EFG1* ORF.** The DNA sequence preceding the ATG start site of the *EFG1* gene in strain ATCC10231 is shown. Sequences marked in red were altered to introduce novel cleavage sites for restriction enzymes (asterisks). These sites and the native *Hpa*I site at position −386 to −391 were used for deletion construction. Transcription start sites are indicated by arrowheads: in the yeast growth form, start sites of the large transcript (3.3 kb) were identified at positions −1170, −1143 and −1112 (amended from Tebarth 2003) and at position −1125 that corresponds to −1117 in strain SC5314 (Bruno 2010). Start sites of a short transcript (2.2 kb) occurring in the yeast form were detected at position −74 (Tebarth 2003) or, in the *opaque* growth form of strain WO-1, at positions −145 and −162 (Srikantha 2000). Sequences representing putative binding sites for the TATA box binding protein (TBP-box) or the Matα2 regulator protein are in green font. A short uORF encoding 4 amino acids, which occurs in strain ATCC10231 but not in strain SC5314, is marked in blue font and is underlined. In addition, putative uORFs starting with a non-ATG sequence (TTG) are marked in blue and green italics, but only uORFs occurring in both ATCC10231 and SC5314 strains are indicated.

**Table S1. Strains**

**Table S2. Oligonucleotides**

